# Major microbiota transitions across liver disease progression are mediated by key taxa linking host environments and microbial communities

**DOI:** 10.64898/2026.07.29.741610

**Authors:** Kenta Suzuki, Tomonori Kamiya, Hideki Fujii, Sotaro Takano, Yoshimi Yukawa-Muto, Norifumi Kawada, Shinji Fukuda, Hiroshi Masuya, Naoko Ohtani

## Abstract

The gut microbiome plays a critical role in chronic liver disease, yet quantitative methods that integrate microbial community structure with clinical indices remain limited. Here, we applied an energy landscape analysis (ELA)-based framework to characterize transitions in gut microbiota along a clinical gradient of liver disease severity, as measured by the FibroScan-AST (FAST) score. The analysis revealed characteristic community states corresponding to low, intermediate, and high FAST scores, indicating that liver disease progression is associated with structured transitions in microbial community organization rather than simple changes in overall diversity. To further dissect community-level organization, we developed the community shaping index (CSI), which quantifies how strongly individual taxa are associated with specific community structures. Integrating CSI with species‒environment association parameters identified key taxa with distinct ecological roles, including unclassified *Subdoligranulum* taxon, an unclassified *Ruminococcus gnavus* group taxon, and *Streptococcus salivarius*, that link host environmental conditions with community composition. Exploratory causal discovery suggested that *Streptococcus salivarius* and the unclassified *Subdoligranulum* may play distinct, potentially active roles in linking liver health with gut microbial organization, but with contrasting associations: *S. salivarius* was linked to disease-related liver deterioration and dysbiotic states, whereas the unclassified *Subdoligranulum* was associated with healthier liver function and coherent community organization. Overall, this ecologically grounded framework provides a unified and mechanistic foundation for analyzing microbiome structure and disease-associated transitions.

## 1. Introduction

Metabolic dysfunction-related steatotic liver disease (MASLD) has emerged as a global health concern, affecting approximately 25% of the adult population worldwide (Younossi et al., 2016). Symptoms may range from simple steatosis to metabolic dysfunction-associated steatohepatitis (MASH), liver fibrosis/cirrhosis, and ultimately hepatocellular carcinoma, representing a continuum of increasing severity. With the global rise in obesity and type 2 diabetes, MASLD is rapidly becoming the leading cause of chronic liver disease. A Delphi consensus process in 2023 led by the American Association for the Study of Liver Diseases and the European Association for the Study of the Liver recommended the name “metabolic dysfunction-related steatotic liver disease” for the group of diseases previously known as non-alcoholic fatty liver disease (NAFLD) (Rinella et al. 2023). This nomenclature reaffirms the importance of both cardiovascular metabolic risk factors and alcohol consumption as factors contributing to disease progression in steatotic liver disease (Rinella et al. 2023, Kanwal et al. 2024). Clinically, MASLD and NAFLD, as well as NASH (non-alcoholic steatohepatitis) and MASH, have been shown to represent essentially the same population, and we therefore use the terms MASLD and MASH throughout this paper (Iwaki et al. 2024, Kamada et al. 2024).

Against the increasing incidence of MASLD, there is a pressing need for accurate, noninvasive methods that enable early detection and effective risk stratification (Loomba & Friedman, 2021). Over the past decade, noninvasive liver disease assessments (NILDA) for MASLD have made significant progress (Sterling et al. 2025a,b). Previously, diagnosis was obtained through liver biopsy, which was invasive and had low accuracy; however, with the advent of various NILDA methods, it is now possible to obtain diagnoses with the same accuracy as liver biopsy. In particular, the FibroScan-AST (FAST) score is a composite index that integrates liver stiffness measurement (LSM) and the controlled attenuation parameter (CAP) from vibration-controlled transient elastography (FibroScan), along with serum aspartate aminotransferase (AST) levels (Newsome et al. 2020). The FAST score is one of the few NILDA currently in use that may reflect not only the progression of liver fibrosis but also disease severity (Newsome et al. 2020, Fujii et al. 2021). Interestingly, it has been also shown to be useful in predicting the onset of hepatocellular carcinoma after the eradication of hepatitis C virus (Ogawa et al. 2020).

Emerging evidence indicates that the gut microbiota is a key modulator of MASLD progression (Aron-Wisnewsky et al. 2020). The gut‒liver axis, established through the portal venous system, enables bidirectional metabolic and immunological communication (Tripathi et al., 2018). Despite growing interest in the role of gut microbiota in MASLD, current approaches to microbiome analysis largely rely on taxonomic profiling, diversity indices, or univariate associations with clinical parameters (Bajaj et al. 2012, 2014; Qin et al., 2014; Lipidot et al., 2020, reviewed by Acharya and Bajaj 2019, Liu et al. 2025, Xirouchakis et al. 2025). These methods lack the capacity to capture the global structural features and stability of microbial communities in relation to continuous disease phenotypes.

To address this gap, we introduce an energy landscape analysis (ELA)-based framework for characterizing how the stability of microbial communities changes along environmental gradients (Suzuki et al., 2021; Fig. 1). The ELA, originally established in the study of neural dynamics (Watanabe et al., 2013), has since gained broader support for its applicability and reliability (Masuda et al. 2025).

**Figure 1.**
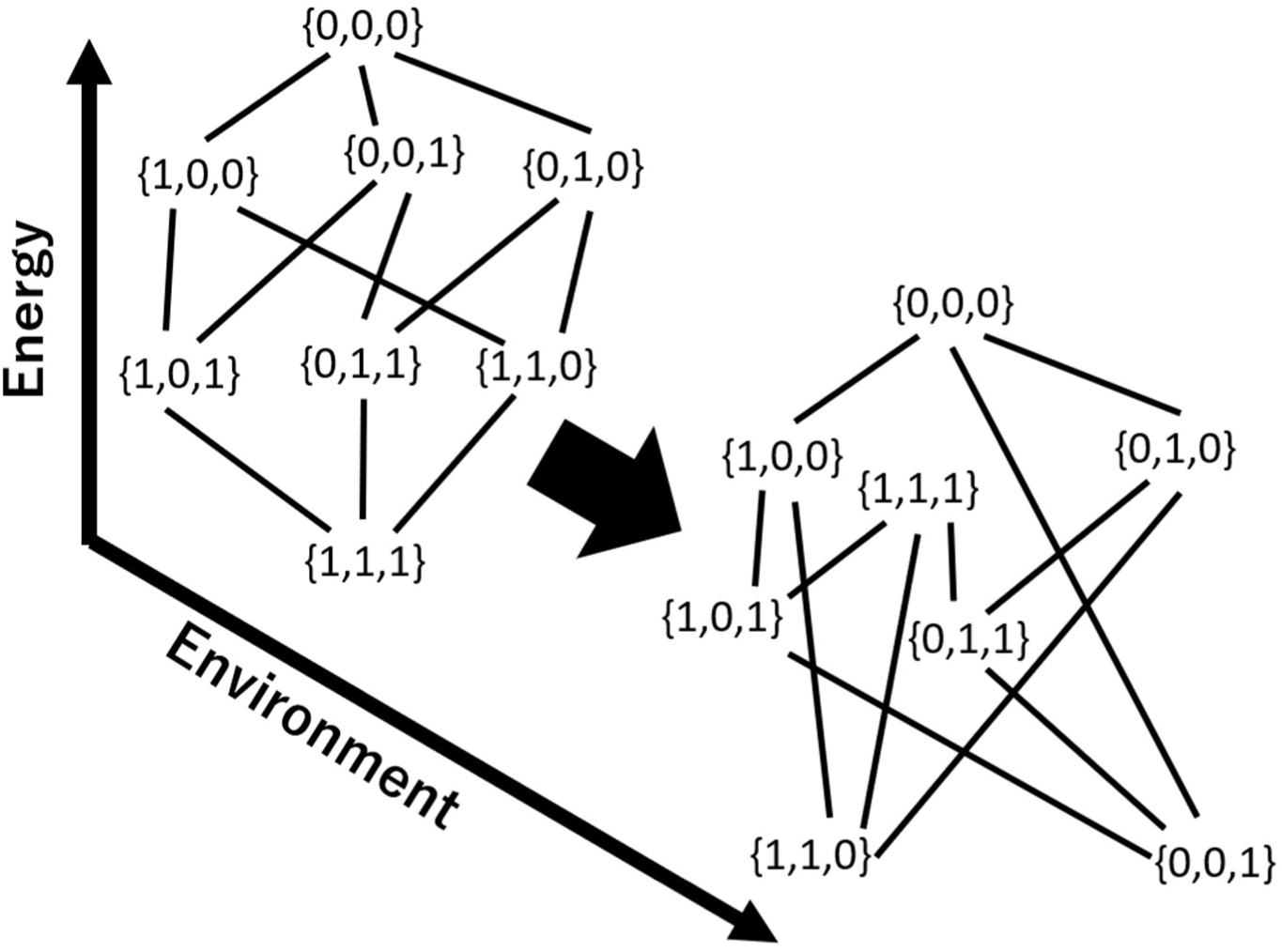
Illustration of microbial stability captured by ELA along an environmental gradient. Along the environmental gradient, the number of energy minima shifts from one ({1,1,1}) to two ({1,1,0} and {0,0,1}), indicating a potential change in the representative community composition. Array of numbers represents presence (1) or absence (0) of the first, second and third species. Energy landscapes allow the stability of a microbial community to be inferred directly from empirical data. Although surface illustrations (ball-and-cup diagrams) are often used as visual metaphor for the concept of a stability landscape, it is important to emphasize that the landscape in this framework is defined on a graph representing different community memberships. For this reason, Suzuki et al. (2021) explicitly distinguish the stability landscape obtained by energy landscape analysis from the traditional ball-and-cup diagrams commonly used in ecology. Conventional stability landscapes represent system dynamics in a continuous state space, such as species abundance or frequency space, as a potential-like surface of the underlying vector field. In such landscapes, valley bottoms correspond to stable equilibria, whereas peaks or ridges separating valleys correspond to basin boundaries, including unstable equilibria. By contrast, the energy landscape in ELA is defined on a discrete space of community memberships, represented as a weighted graph in which each node is a presence‒absence configuration. Thus, in ELA, local energy minima and basin boundaries are interpreted within this discrete membership space rather than in a continuous abundance or frequency space. The landscape therefore reflects the probability of observing each community-membership configuration after the effects of the underlying ecological dynamics are considered in aggregate. Moreover, although the method is derived from model equations (1-3), it essentially provides a weighted representation of the membership and is therefore fundamentally model-free. As a result, it can be applied independently of the specific dynamical equations or the physical system under consideration.

Emerging work in both ecology (Suzuki et al. 2021; Fujita et al., 2023; Miyamoto et al., 2023; Fujita et al., 2025; Kadoya et al., 2025; reviewed by Toju et al. 2026) and medical data analysis (Yamamoto et al., 2024; Ito et al., 2025; Tatematsu et al. 2026) demonstrates a strong ability to infer dynamical properties of complex empirical systems from cross-sectional data, extending beyond applications in neuroscience. We reconstructed energy landscapes by integrating taxon‒environment and taxon‒taxon relationships, enabling the decomposition of both types of associations. Unlike conventional statistical approaches, this approach identifies distinct microbiome states that reflect underlying ecological processes captured by an explicit model. Building on this unified model, we introduce the community shaping index (CSI), which quantifies the extent to which individual taxa are associated with specific community structures. By integrating this index with inferred taxa‒environment relationships and assessing the causal directionality of these links through statistical causal inference, we demonstrate how individual taxa mediate the association between environmental factors and community-level organization. Previous applications of ELA provided valuable insights into stability and state transitions in various empirical systems (Masuda et al., 2025; Toju et al. 2026). To our knowledge, this study is the first to extend ELA from identifying community-state transitions along environmental gradients to quantifying how individual taxa link host environmental conditions with community-level organization, by integrating taxon-specific environmental responses, taxon‒taxon associations, CSI, and exploratory causal inference.

## 2. Methods

### 2.1 Study participants, metadata, and screening

This study was approved by the Ethics Committee of Osaka City University (now known as Osaka Metropolitan University) School of Medicine (approval number: 3722), and written informed consent was obtained from all adult participants. All methods were carried out in accordance with relevant guidelines and regulation.

For each participant, fecal samples were collected for 16S rRNA gene sequencing analysis, along with clinical and physiological metadata, including age, sex, weight, height, and body mass index (BMI), which are summarized in Table 1. Liver disease status was assessed by the FAST (FibroScan-AST) score, calculated by the following logistic regression-based formula (Newsome et al., 2020):

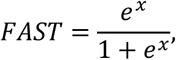

**Table 1.**
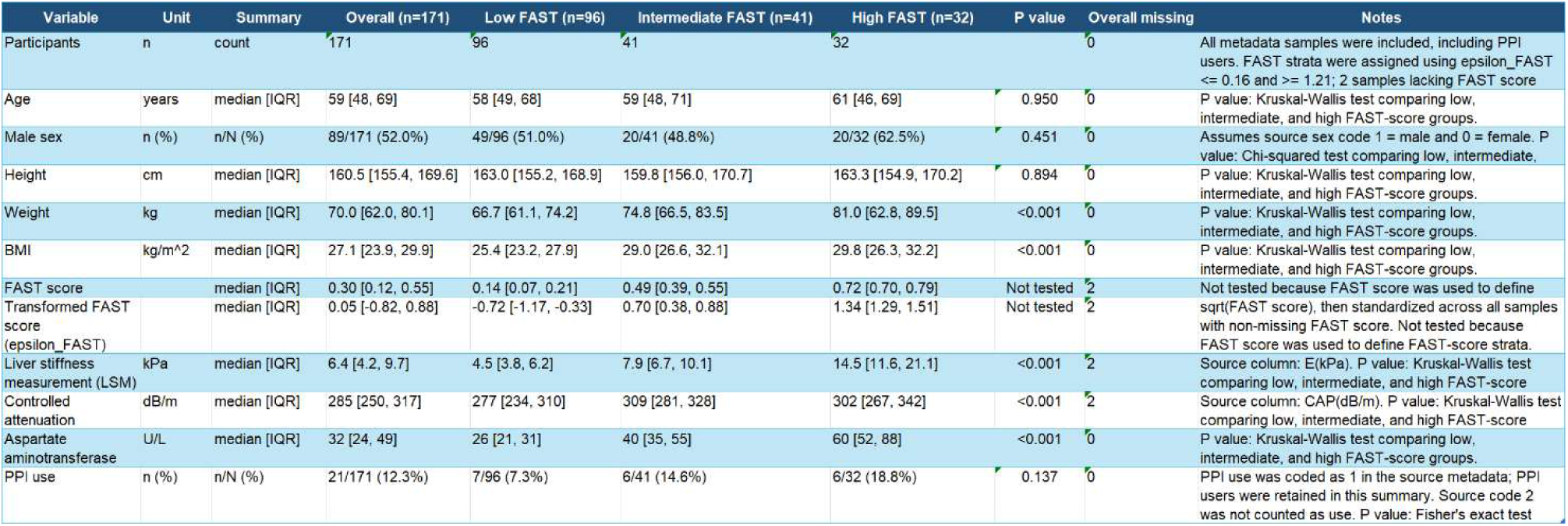
Demographic and clinical characteristics of the study participants. Demographic and clinical variables were summarized for all 171 enrolled participants, including proton pump inhibitor users. Participants were stratified into low, intermediate, and high FAST-score groups based on transformed FAST score values. Samples with missing FAST score values were included in the overall cohort but excluded from FAST-score stratified summaries. Continuous variables are shown as mean ± standard deviation, and categorical variables are shown as counts and percentages. P values compare low, intermediate, and high FAST-score groups. Continuous variables were compared using the Kruskal‒Wallis test, and categorical variables were compared using the chi-squared test or Fisher’s exact test when appropriate. FAST score was not tested because it was used to define the strata.

where

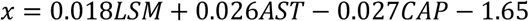

Here, liver stiffness measurement (LSM) is expressed in kPa, controlled attenuation parameter (CAP) in dB/m, and serum aspartate aminotransferase (AST) in U/L. In addition to liver-related clinical indicators, participants provided information on proton pump inhibitors (PPI)usage (see Supplementary Table S1 for the FAST score and PPI value of each sample).

To reduce skewness and align the scale with other terms in the equation, the FAST score was subjected to square root transformation and rescaled to have a mean of 0 and standard deviation of 1. The transformed variable is denoted as ε_*FAST*_. Based on previous clinical validation, we used FAST score cutoffs of 0.35 and 0.67 as reference values for clinically low and high FAST-score ranges, respectively, rather than as definitions derived from our microbiome-based energy landscape analysis. A FAST score of 0.67 (=ε_FAST_ = 1.21) has been reported to have a positive predictive value of 0.69 for NAFLD activity score ≥4 and fibrosis stage ≥2, whereas a FAST score of 0.35 (ε_*FAST*_ = 0.12) has been reported to have a negative predictive value of 0.94 for the same condition (Newsome et al. 2020). Accordingly, values between these two thresholds were regarded as corresponding to a clinically intermediate FAST-score range. We then compared these clinically defined reference ranges with the low-, intermediate-, and high-FAST community states identified by the energy landscape analysis.

### 2.2 Sample collection, DNA extraction, and 16S rRNA gene sequencing and processing

Fresh human fecal samples were obtained using stool collection tubes (Techno Suruga Laboratory CO., Ltd., Shizuoka, Japan). Fecal DNA was extracted by BIKEN Biomics Inc. (Osaka, Japan) using an automated DNA extraction machine (GENE PREP STAR PI-480, Kurabo Industries Ltd, Osaka, Japan) in accordance with the manufacturer’s protocol with modifications as needed to optimize yield and purity.

The V1‒V2 region of the bacterial 16S rRNA gene was amplified using universal primers 27Fmod (5’- AGRGTTTGATCMTGGCTCAG-3’) and 338R (5’-TGCTGCCTCCCGTAGGAGT-3’). Amplicons were sequenced on a MiSeq system (Illumina) using a MiSeq Reagent v2 500 cycle kit with 251-bp paired-end reads. Raw sequences were quality filtered, trimmed, denoised, and merged using the Quantitative Insights into Microbial Ecology 2 version 2019.4 (QIIME2) pipeline and DADA2 workflow. Amplicon sequence variants (ASVs) were taxonomically assigned using the SILVA v132 reference database.

### 2.3 Data preprocessing

Of the 171 participants enrolled, 170 had microbial abundance profiles and associated metadata. To ensure data quality and consistency, we excluded individuals lacking FAST score records, those using proton pump inhibitors, and those with fecal samples with atypical sequencing depth. Specifically, sequencing read count profiles were assessed using Gaussian mixture model (GMM)-based clustering, and samples belonging to the largest GMM-derived cluster were retained as the primary high-quality dataset (Supplementary Figure S1 and Supplementary Table S1). After application of these criteria, data from 104 participants were included in the final analysis. Of these, 19.8% and 55.6% of the cohort exhibited high (ε_*FAST*_ ≥ 1.21) and low (ε_*FAST*_ ≤ 0.16) FAST scores, respectively.

### 2.3 Energy landscape analysis

In ELA, the community composition σ is defined as a binary vector of length *S*, where *S* is the total number of taxa. Therefore, there is a total of 2*^S^* unique community compositions, which are the nodes of the alternate state. We denote a *k*th community composition (*k* ∈ {0,1, …, 2^*S*–1^}) as σ^(*k*)^ = 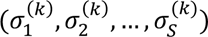, where 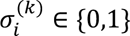 is the presence/absence status of the *i*th taxon. Next, the links that connect the community compositions (i.e., σ values) are defined under the assumption that community compositions change in a stepwise manner; that is, two community compositions are joined by a link only if they have the opposite status (i.e., 0/1) for just one taxon. Under this assumption, the community compositions form a regular network in which each node has *S* links.

To construct an environmentally extended energy landscape, we first define the energy of community compositions (i.e., nodes of the network) based on environmental conditions and organismal occurrence patterns, using the extended maximum entropy model (Suzuki et al. 2021). Generally, the model determines the probability of the occurrence of a community composition σ^(*k*)^ under a given environmental condition defined as a vector of *M* environmental variables, s = (ε_1_, ε_2_, … , ε_*M*_). The probabilities of community states *P*(σ^(*k*)^|s) at environment s are given by

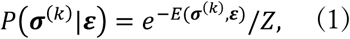

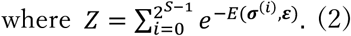

Here, the energy *E* for the community state σ^(*k*)^ at environment s is

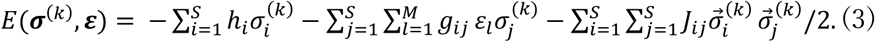

In this model, 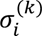 represents the presence/absence status of species *i* in *k*th community composition, *ε_l_* represents *l*th environmental factor, ℎ_*i*_ represents the net effect of implicit abiotic factors by which the *i*th taxon is more likely to be present (ℎ_*i*_ > 0) or not (ℎ_*i*_ < 0), *g*_*lj*_ represents the effect of the *l*th environmental factor on the *j*th taxon, and *J_ij_* represents taxon‒taxon association based on a co-occurrence pattern of the *i*th and *j*th taxon. *E*(σ^(*k*)^, ε) is referred to as “energy” following the terminology of statistical physics. The difference in the energy of two adjacent community compositions represents the directionality of state transitions in the sense that a lower energy state is more likely to be realized than a higher energy state.

In our analysis, s consists of a single variable that represents the transformed FAST score (ε_*FAST*_). Thus, specifically for our analysis, equation (3) is

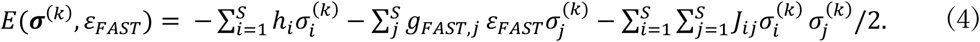

Here, *g_FAST,j_* represents the effect FAST score on the *j*th taxon. Because the logarithm of the probability of a community state is inversely proportional to *E*(σ^(*k*)^, ε_*FAST*_), a community state having lower *E* is more frequently observed with the given ε_*FAST*_ value.

The parameters of the extended pairwise maximum entropy model in equation (4) were adjusted to the occurrence table and environmental factor (ε_*FAST*_) using the *runSA* function in the rELA software package (https://github.com/kecosz/FASTexELA) after the hyper parameters (length of optimization steps and regularization coefficients) were selected using the *Findbp* function. In this analysis, the occurrence table was constructed from the sample-wise read count table as follows: taxa with relative abundance ≥1% were considered “present” (1); those below this threshold were considered “absent” (0). Taxa with presence in ≤5% (negligible influence) or ≥95% (almost always present) of the samples were excluded from the analysis. As a result, 58 taxa were retained for downstream analysis. The total relative abundance of these taxa across all samples accounted for a mean of 68.5% and a standard deviation of 10%.

The maximum likelihood estimates of ℎ*_i_*, *g_ij_*, and *J_ij_* were obtained by a stochastic approximation method (Suzuki et al. 2021). We focused on *g_FAST,i_* and *J_ij_* as key estimates in our analysis. Specifically, *g_FAST,i_* was interpreted as the preference of each taxon for the FAST score, where taxa with higher *g_FAST,i_* values tend to appear more frequently in high FAST score environments; and *J_ij_* was used as an indicator of biotic interactions.

Once these parameters were estimated, equation (4) assigned an energy value to every possible community composition at each value of ε_FAST_. The regular graph of community compositions was therefore represented as a weighted graph, in which each node corresponded to a presence‒absence configuration and its node weight was given by *E*(*σ^(*k*)^*, ε_FAST_). On this graph, an energy minimum was defined as a community composition whose energy was lower than that of all directly connected neighboring compositions. Following Suzuki et al. (2021), these energy minima were interpreted as *stable states* in ELA and are referred to here as representative community compositions.

Energy minima were identified using a Monte Carlo-based gradient-descent procedure implemented in the *gradELA* function of the rELA package. For each value of ε*_FAST_*, 20,000 initial community compositions were randomly sampled. Starting from each initial composition, the algorithm repeatedly moved to the directly connected neighboring composition with the largest decrease in energy until no neighboring composition had a lower energy value. The terminal composition reached by this procedure was recorded as an energy minimum. The basin of an energy minimum was defined as the set of community compositions that converged to the same energy minimum under this gradient-descent procedure. Adjacent community compositions belonging to different basins were regarded as forming basin boundaries.

We applied this procedure across different ε_FAST_values from -2 to 2 at intervals of 0.1 using the *gradELA* function and visualized the resulting changes in energy minima and their basins as a stable-state diagram using the *showSSD* function.

### 2.4 FAST score prediction by environmentally-extended ELA

Whether a given community composition is better adapted to high or low FAST score environments was quantified by the following measure:

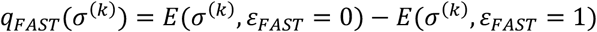

If this value is substantially positive, i.e., *E*(*σ^(*k*)^*, ε_*FAST*_ = 0) > *E*(*σ^(*k*)^*, ε_*FAST*_ = 1), it means that the community is more likely to be observed under high ε_*FAST*_ conditions. Conversely, if this value is negative, i.e., *E*(*σ^(*k*)^*, ε_FAST_ = 0) < *E*(*σ^(*k*)^*, ε_*FAST*_ = 1), it indicates that the community is more prevalent in low ε_*FAST*_ environments. Although this definition is mathematically very simple and could be further generalized, it is sufficient for the present study because environmental dependence is modeled linearly throughout our analysis.

In this study, we adopted *q*_*FAST*_ as an index for predicting the FAST score from microbial composition within the ELA framework. It is evident from equation (4) that

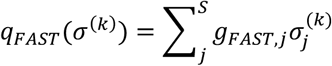

This means that the magnitude of *q*_*FAST*_ is ultimately determined by the sum of *g*_*FAST*_ values for taxa present in *σ^(*k*)^* (i.e., 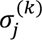 = 1). Communities that tend to appear under high FAST score conditions are composed predominantly of taxa with large positive *g*_*FAST*_ values, if other conditions are the same.

To ensure robust performance estimates across different data splits, we randomly shuffled the data 20 times and performed 10-fold cross-validation for each iteration. Because *q*_*FAST*_ does not aim to predict the actual FAST score values directly, predictive performance was evaluated by calculating the Spearman rank correlation coefficient between *q*_*FAST*_ and ε*_FAST_*. This evaluation procedure was consistently applied across all machine learning methods explained in the next section and the correlation-based (COR) method described below.

For comparison with *q_FAST_*, we also employed a simpler method in which the Spearman correlation coefficient between each taxon’s relative abundance and ε_FAST_ (*ρ*_*FAST*_) was computed. The sum of *ρ*_*FAST*_ values for all taxa present in a sample was then used as the predictive score. These correlations were calculated on the raw relative abundance values, prior to their conversion into binary presence/absence. This method is referred to hereafter as the COR method.

### 2.5 Machine learning methods

Three types of regression models—linear regression (LR), support vector regression (SVR) and random forest (RF) —were used to predict continuous environmental variables from microbiome data, in addition to ELA and the COR method. The R packages, functions, and primary parameter settings used for each model are described below.

The LR model was constructed using the *lm* function in the *stats* package in R. The response variable was ε*_FAST_*, with all microbial relative abundance as predictors. No additional parameter tuning, regularization, or feature selection was applied. Support vector regression (SVR) was performed using the *svm* function in the *e1071* package (Dimitriadou et al. 2009) with *type = “eps-regression”* and otherwise default settings, including the default radial basis function kernel and default hyperparameter. Random forest modeling was implemented using the *ranger* function in the *ranger* package (Wright and Ziegler 2017). The model was used in default regression mode without any manual tuning of hyperparameters such as *num.trees* or *mtry*. As with the COR method, this machine learning task used the relative abundance data before their conversion to presence/absence. These models were included as standard methodological benchmarks rather than as fully optimized predictive models.

### 2.6 Community shaping index (CSI)

To quantify the contribution of each taxon to the different community states, we computed a community shaping index (CSI) for a community composition *k* based on the pairwise interaction matrix *J* and the membership of the community *σ^(*k*)^*. For taxa *i* in community *k*, it is,

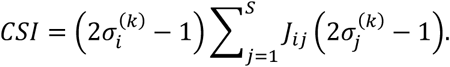

If taxon *i* is a member of community *k*, positive *J_ij_* values with other members of *k* increase the CSI, while negative *J_ij_* values decrease it. If *i* is not included in *k*, then positive *J_ij_* values with other taxa that are also absent from *k* increase the CSI, and negative *J_ij_* values decrease it; however, this “absence-based” effect on community composition is not the focus here. In short, the CSI represents the total contribution of taxon *i* to community *k*, incorporating both its positive associations within the community and its suppressive effects on alternative community compositions.

A general interpretation of the CSI can be summarized as follows. A taxon has a high CSI for a given community composition when it has positive associations with members of that community and, conversely, negative associations with taxa outside that community. Thus, a large positive CSI indicates that the taxon tends to be strongly associated with that community composition. However, this should not be interpreted as evidence of a causal effect. Such an association may include both active cases, in which the taxon contributes to the formation of the community composition, and passive cases, in which the occurrence of the taxon is promoted by that composition. Therefore, to further examine the causal directionality of the relationships between taxa and community structures suggested by high CSI values, we introduced an additional causal inference analysis.

### 2.7 Causal inference

To explore potential causal relationships among liver condition, a microbial taxon, and global gut community structure, we conducted causal discovery using the Fast Adjacency Skewness (FASK) algorithm (Sanchez-Romeo et al. 2019). We constructed a data table consisting of three variables: (i) liver health status quantified by FAST score; (ii) the relative abundance of focal taxa; and (iii) a community composition indicator (community membership score; CMS) representing the proximity of each sample to a specific representative community composition (energy minimum) on the energy landscape. Therefore, this causal index is calculated for each combination of representative community composition *C* (or the union of compositions when *C* represents a group of multiple compositions), focal taxon, and FAST score. All variables were treated as continuous and standardized prior to analysis. Microbial abundance data were preprocessed to account for the compositional nature of microbiome data. Specifically, relative abundances were transformed using a centered log-ratio transformation after adding a small pseudocount to handle zeros.

FASK is a score-based method that infers causal structure by exploiting non-Gaussianity (skewness) in the data distribution. FASK can be applied to cross-sectional datasets and allows for cyclic causal relationships, making it suitable for biological systems where feedback mechanisms may exist. The algorithm combines adjacency search based on a score criterion with skewness-based orientation rules to infer directed and potentially bidirectional causal effects. Here, adjacency search was conducted using the SEM-BIC score, with the penalty discount set to 2.0 to control model complexity. Edge orientation was performed using the FASK v2 left‒right rule, with the skewness threshold for additional edge inclusion set to 0.05 and the orientation bias parameter (*faskDelta*) set to 0.2. The maximum conditioning set size was unrestricted (depth = –1), and resampling procedures were not applied.

The CMS was calculated as follows. For a target community, the index was computed by aggregating the relative abundances of taxa belonging to that community and normalizing this value by the total microbial abundance in the sample. Importantly, when evaluating causal relationships involving a specific taxon, that taxon’s abundance was excluded from the community index calculation to avoid circular dependence. Thus, the community index captures a global property of community organization that is conceptually distinct from the abundance of any single taxon.

### 2.8 Statistical methods

We evaluated microbial alpha diversity using four indices—species richness (number of observed taxa per sample), Shannon entropy, Simpson’s index, and Pielou’s evenness—and examined their correlations with the FAST score. Shannon entropy and Simpson’s index were calculated using the *diversity* function in the R library *vegan* (CRAN), with the index parameter set to *shannon* and *simpson*, respectively. Pielou’s evenness was computed by dividing the Shannon entropy by the logarithm of species richness, in accordance with its definition. To evaluate correlations between taxon abundances while accounting for the compositional nature of the data, we employed SparCC (Friedman and Alm 2012), implemented via the SpiecEasi library (Kurtz et al. 2015). For testing the similarity between matrices, we used the *mantel* function in *vegan*. Lastly, we conducted permutational multivariate ANOVA (PERMANOVA) using the *adonis2* function in *vegan* to statistically assess the influence of the continuous variable on community composition.

We performed a sensitivity analysis of the entire analytical workflow to assess the robustness of the results. Specifically, within a range that included the main analysis setting, we tested five relative-abundance thresholds for constructing microbial taxon presence‒absence matrices: 0.005, 0.008, 0.010, 0.012, and 0.015. We also tested five combinations of lower and upper occurrence-frequency thresholds for retaining taxa: 0.01‒0.99, 0.03‒0.97, 0.05‒0.95, 0.07‒0.93, and 0.10‒0.90. Therefore, excluding the main setting, there were 24 candidate parameter combinations. However, the analysis was restricted to conditions in which the number of retained taxa was within ±25% of that in the main analysis, namely datasets in which 44 to 72 taxa were selected, relative to the 58 taxa retained in the main analysis (Supplementary Table S2). We considered this restriction reasonable to maintain a balance between sample size and the number of variables, as well as consistency of interpretation. The parameters estimated under the 18 selected conditions were broadly consistent with those obtained in the main analysis (Supplementary Figure S2, S3, S4). We therefore conducted further analyses to compare these results with the main analysis and evaluated whether the identification of key taxa and the interpretation of community states were robust to differences in preprocessing conditions.

### 2.9 Code and Data availability

The datasets generated and/or analyzed during the current study are available in the DDBJ database (https://ddbj.nig.ac.jp/search) under accession number PRJDB40175. A self-contained package containing the code and datasets required to reproduce all analyses is available in the GitHub repository: https://github.com/kecosz/FASTexELA.

## 3. Results

### 3.1 Microbiome analysis incorporating environmentally-extended ELA

To verify the reliability of the model acquired, we compared its predictive performance on FAST scores with that of other representative machine learning methods (linear regression [LR], support vector machine [SVM], and random forest [RF]), as well as predictions based on correlation coefficients (COR). Even though the other methods utilized relative abundance data, the predictions by environmentally-extended ELA outperformed them all (Fig. 2; Supplementary Figure S5). This outcome both supports the validity of our approach, which explains community occurrence probabilities through environment-taxa and taxon‒taxon relationships, even compared to more flexible models, and reinforces the appropriateness of interpreting microbial communities through its estimated parameters.

**Figure 2.**
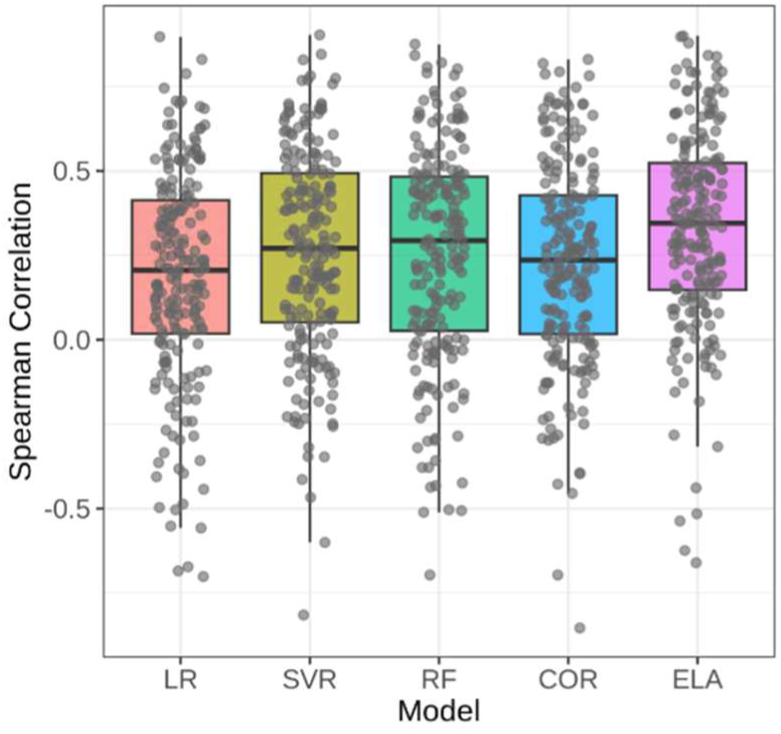
Comparison of FAST score prediction by statistical or machine learning methods (linear regression [LR]; support vector regression [SVR], random forest [RF]), environmentally-extended energy landscape analysis (ELA), and correlation-based (COR) method. Whiskers indicate minimum and maximum values; lower and upper edges of the boxes correspond to 25th and 75th percentiles, respectively; horizontal lines inside the boxes indicate the median.

In terms of the overall gut community structure, our approach revealed that representative community compositions (stable states) associated with low FAST scores exhibited increased energetic instability and were subsequently replaced by compositions dominated by taxa associated with high FAST scores, which then showed greater energetic stability (Fig. 3a-c). This observation indicates that variation in FAST scores is closely aligned with shifts in gut microbiota composition. Based on this and the hierarchical clustering of community compositions (Fig. 3d), we grouped the representative compositions by compositional similarity, not by predefined clinical FAST-score thresholds. We then interpreted these composition-based groups according to their locations along the ε_FAST_ gradient as low-, intermediate-, and high-FAST-associated community states: (C1, C2, C3), (C4, C5.1‒3, C6), and (C5.4, C5.5, C8), respectively. Although C7 appeared at intermediate ε_*FAST*_ values, its community composition was close to those of C5.4, C5.5 and C8, suggesting that it may represent a somewhat exceptional composition. Within the FAST score range where each stable state exists, there was at least one observed community composition that belonged to the basin of that stable state (Fig. 3e). Such stepwise shifts were common across all sensitivity analyses, despite differences in the compositions of the stable states (Supplementary Figure S6, S7).

**Figure 3.**
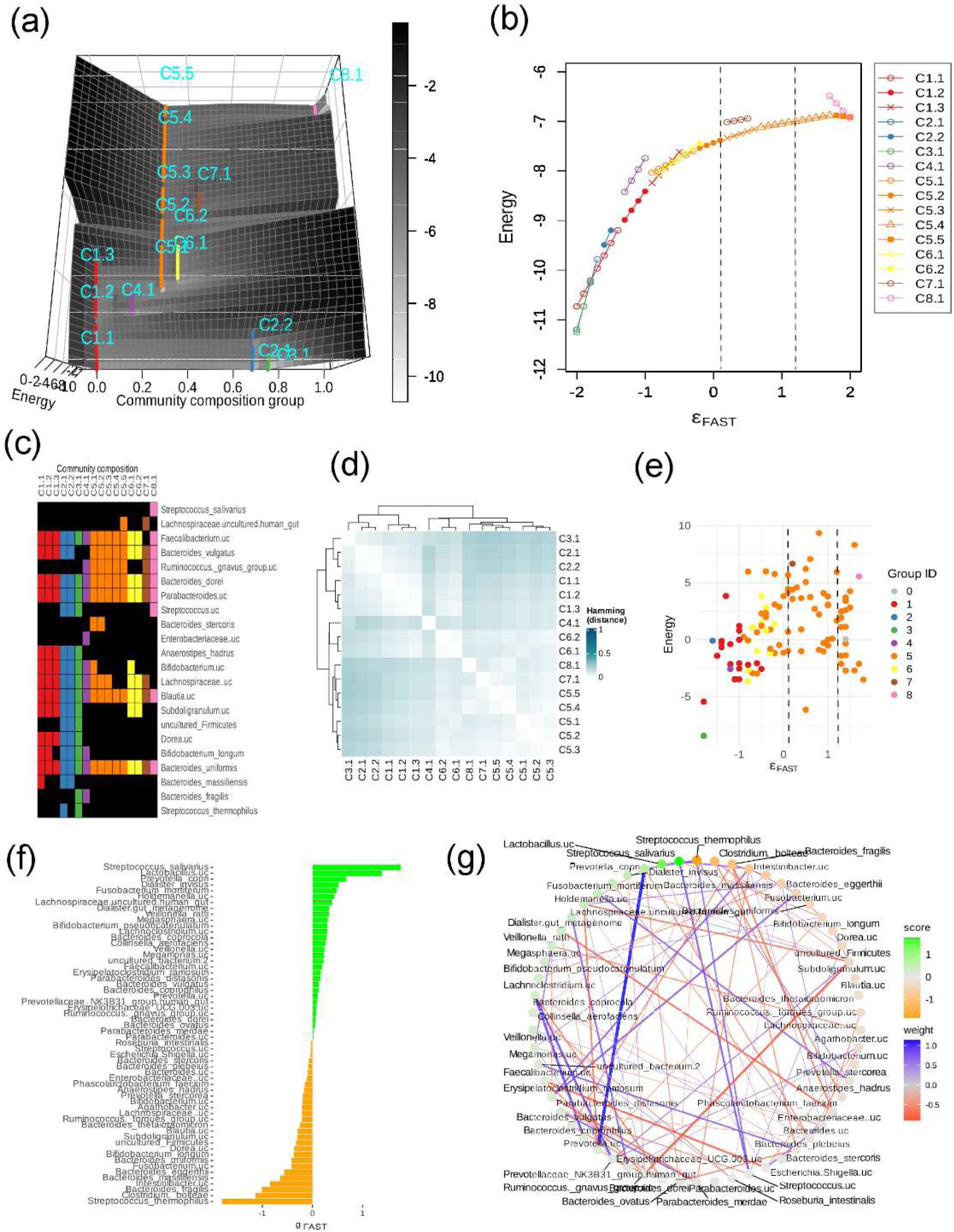
Microbiota structure associated with FAST score. (a) Three-dimensional surface plot of energy landscape parameterized by ε*_FAST_*. Points on the landscape correspond to energy minima. Different colors represent distinct groups of community compositions, and each symbol type indicates an identical community composition. See panel (c) for the correspondence between community composition colors and indices. (b) Stable-state diagram. The y-axis plots the energy of energy minima in the energy landscape and the x-axis plots ε*_FAST_*; the range of ε_*FAST*_ in which a stable state within a group of community composition is represented as points and line segments. We grouped community compositions in accordance with two criteria: (i) under minimal environmental changes, adjacent compositions differed by no more than three species; and (ii) the energy difference between adjacent compositions was less than 1.2 times the maximum within-group energy variation. The dashed lines indicate 0.12 and 1.2, which were used as clinically validated thresholds separating low, intermediate and high FAST scores. Please note that this does not necessarily correspond to the high, intermediate or low FAST-score ranges associated with the occurrence patterns of community compositions discussed in the main text. (c) Stable-state community compositions visualized as tile plots. Only taxa included in any of the community states are shown, ordered by descending *g_FAST_*. (d) Heat map of pairwise distances between community compositions (normalized Hamming distances) and the results of hierarchical clustering based on these distances. (e) Plot of calculated energy values based on community membership (y-axis) versus corresponding FAST score (x-axis) for observed community compositions. Colors indicate basin membership (i.e., the basin of energy minimum to which the community belongs). The dashed lines indicate 0.12 and 1.2, which were used as clinically validated thresholds separating low, intermediate and high FAST scores. (f) *g*_*FAST*_ values for each taxon. Higher values indicate stronger associations with high-FAST environments; lower values reflect associations with low-FAST environments. (g) Network based on estimated taxon‒taxon interactions (*J_ij_*). Blue edges represent positive associations (tendency to co-occur); red edges represent negative associations (mutual exclusivity). Color intensity and width of links reflect the strength of the interaction. Node colors correspond to *g*_*FAST*_ values.

Focusing on the representative community compositions captured by the energy landscape, our results clearly showed that several taxa observed in samples with low FAST scores were lost at intermediate to high FAST scores (Fig. 3a‒c). For example, *Anaerostipes hadrus*, *Bacteroides massiliensis*, and unclassified taxa assigned to *Subdoligranulum*, *Firmicutes* and *Dorea* appeared only in C1‒C3. These taxa generally had low *g*_FAST_values, consistent with their preferential occurrence in samples with low FAST scores. In contrast, an unclassified *Ruminococcus gnavus* group taxon was broadly observed in the intermediate- to high-FAST compositions C4‒C8. In addition, an unclassified taxon in the Lachnospiraceae family (*Lachnospiraceae uncultured human gut*) and *Streptococcus salivarius* were characteristic of the high-FAST community compositions, particularly C5.5 and C8. Notably, *Streptococcus salivarius* showed the highest *g*_*FAST*_ value, consistent with this trend, whereas the *g*_*FAST*_ values of the *R. gnavus* group taxon were not necessarily high (Fig. 3f). This point is examined in more detail in the following analyses. In the sensitivity analysis, *Anaerostipes hadrus*, *Bacteroides massiliensis*, *Streptococcus salivarius*, and unclassified taxa assigned to *Subdoligranulum*, Firmicutes, *Dorea*, the *Ruminococcus gnavus* group, and Lachnospiraceae were included in all datasets (Supplementary Table S4). However, the proportions of datasets in which at least one stable state contained these taxa were 73%, 100%, 53%, 79%, 58%, 79%, 95%, and 84%, respectively (Supplementary Table S5). In particular, *Streptococcus salivarius* was not included in stable states when the binarization threshold was high (0.015) or when the taxon-retention criterion was stringent (0.10‒0.90). This may partly reflect the relatively small number of patients with severe liver disease in the dataset. Overall, however, these taxa showed consistent general tendencies in their occurrence along the FAST score gradient, as described above (Supplementary Table S5).

Although community-level differences were aligned with FAST scores, α-diversity indices—including species richness, Simpson’s diversity, Shannon entropy, and Pielou’s evenness—did not show significant correlations with FAST scores (Supplementary Figure S8; Supplementary Table S6).

PERMANOVA indicated that only ∼1.4% of variance could be explained by the FAST score (Supplementary Table S7). Therefore, conventional evaluations on overall diversity were unable to capture the relationship between the community structure and FAST score. The correlation between *g*_*FAST*_ and *ρ*_*FAST*_ (defined as the Spearman correlation between relative abundance and FAST score across samples) was 0.78 (Supplementary Figure S9). However, because correlation coefficients are influenced by interspecific relationships, *ρ*_*FAST*_ alone does not fully account for the magnitude of *g_FAST_*. Similarly, across the sensitivity analyses, none of the diversity indices showed a significant correlation with the FAST score, and the proportion of variance explained by the FAST score in PERMANOVA was also small (Supplementary Table S6, S7).

Taxon‒taxon relationships also exhibited complexity but followed certain patterns (Fig. 3g). No significant similarity was observed between the inferred interaction matrix and the SparCC-based correlation matrix (Supplementary Table S8). Therefore, J is considered to capture taxon-taxon relationships that differ from simple correlations.

### 3.2 Link between environmental response and species relationship

For each taxon, we calculated the community shaping index (CSI) for each community composition using the taxon-taxon relationship (Supplementary Figure S10) and plotted it for communities (C1‒C8; Fig. 4). To investigate the directionality of causal relationships among the FAST score, the abundance of individual taxa, and the community membership score (CMS), we further applied FASK, a statistical causal discovery method, which represents degree of bias toward characteristic community compositions evaluated after excluding the abundance of the focal taxon (Supplementary Table S9).

**Figure 4.**
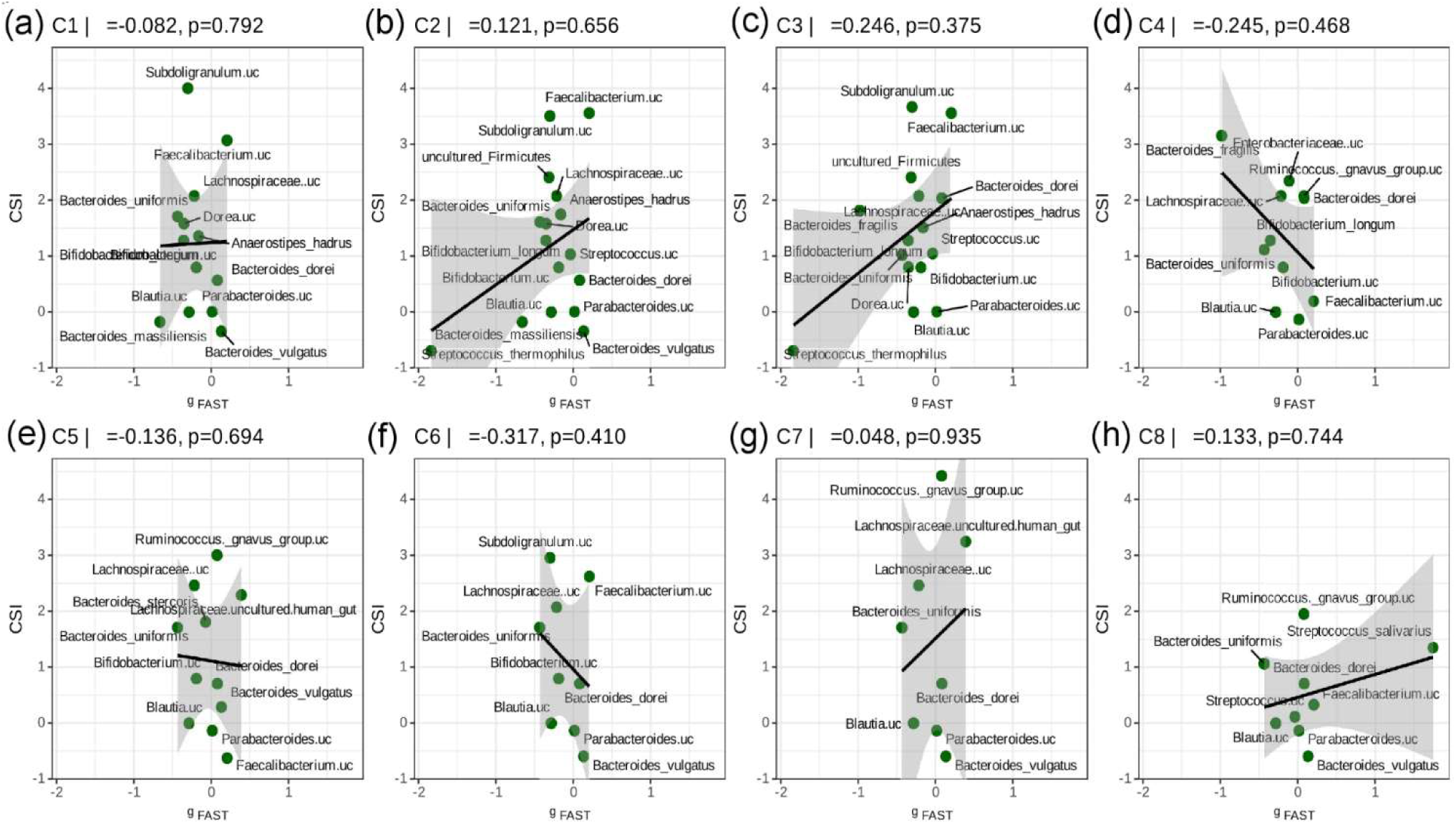
Taxa‒environment relationship (*g*_*FAST*_) and community shaping index (CSI) for C1‒C8. Panels (a‒f) correspond to community compositions C1‒C8. Numbers above each panel indicate Spearman’s rank correlation coefficients and corresponding *p*-values. In these plots, *g*_*FAST*_ represents the taxon-specific association with the FAST-score gradient, whereas CSI represents the strength of association between each taxon and the corresponding community composition. Taxa with large absolute *g*_*FAST*_ values and high positive CSI values can be interpreted as taxa that both respond strongly to the FAST-score environment and are strongly associated with the focal community composition. Accordingly, taxa in the upper right region are linked to high FAST scores and the focal community composition, whereas taxa in the upper left region are linked to low FAST scores and the focal community composition.

Overall, taxa with *g*_*FAST*_ values close to zero, which are relatively insensitive to the FAST score, tended to show large variation in CSI, and no significant correlation was observed between *g*_*FAST*_ and CSI (Fig. 4). This suggests that taxa more strongly associated with the FAST-score environment are not necessarily those that contribute most strongly to shaping the corresponding community composition.

In the representative compositions associated with low FAST scores (C1, C2, and C3), the unclassified *Subdoligranulum* and *Faecalibacterium* showed high CSI values (Fig. 4a‒c). In particular, the unclassified *Subdoligranulum*, a taxon characteristic of C1, C2, and C3, was placed by the exploratory FASK analysis upstream of both the FAST score and community composition in the inferred graphs for all three compositions (Supplementary Table S9). This result suggests that the unclassified *Subdoligranulum* may be involved in linking liver health with low-FAST-associated community organization. In the representative compositions associated with intermediate-to-high FAST scores (C5, C7, and C8), the *R. gnavus* group taxon showed the highest CSI values (Fig. 4e, g, h). Interestingly, a strong mutually exclusive relationship was observed between the unclassified *R. gnavus group taxon* and the unclassified *Subdoligranulum* (Fig. 3g). These two taxa may therefore play contrasting roles in separating community compositions associated with low FAST scores from those associated with intermediate-to-high FAST scores. However, the FASK-based causal analysis did not indicate a consistent relationship between the *R. gnavus* group taxon and either the FAST score or community composition (Supplementary Table S9).

Finally, we note the role of *Streptococcus salivarius*. *S. salivarius* had the highest *g*_*FAST*_ value (Fig. 3f) and the second-highest CSI in C8, the composition associated with the highest FAST scores (Fig. 4h). Moreover, the exploratory FASK analysis placed *S. salivarius* upstream of both the FAST score and community composition in the inferred graph (Supplementary Table S9). Together, these findings suggest that *S. salivarius* may play a potentially active role in linking liver disease severity with high-FAST-associated dysbiotic community organization. As discussed in more detail below, this is a suggestive finding, given that this taxon has been associated with more severe liver disease states, including hepatocellular carcinoma.

Across all sensitivity analyses, the correlation coefficients of *g*_*FAST*_ (Supplementary Figure S4) and CSI values (Supplementary Figure S6) with those from the main analysis were sufficiently high, ranging from 0.910 to 0.999 and from 0.556 to 0.972, respectively. The values for the taxa described above also showed substantial consistency. In the taxon‒taxon relationships represented by *J*, a negative association between the *R. gnavus* group taxon and the unclassified *Subdoligranulum* was also a consistent trend (Supplementary Figure S3). Therefore, the results obtained here can be regarded as robust to differences in binarization thresholds and taxon-selection criteria. Because FASK analyzes continuous variables, it was not included in this sensitivity analysis.

## 4. Discussion

Understanding how gut microbiota structure relates to host phenotypes, particularly in complex clinical settings such as liver pathology, remains a central challenge in microbiome research. Here we addressed this issue by using environmentally-extended form of ELA (Suzuki et al. 2021), a modeling approach that integrates ecological processes with empirical data, to characterize the continuous relationship between microbiota composition and FAST score. A major finding of our analysis is that liver disease severity, as represented by the FAST score, was not simply associated with gradual changes in individual taxa, but with transitions among representative community compositions. These community-level changes were captured as shifts in energy minima along the FAST-score gradient, suggesting that gut microbial organization may undergo structured state transitions during liver disease progression.

This community-level perspective is important because conventional summary measures did not sufficiently explain the observed relationship between the microbiota and FAST score. Alpha-diversity indices, including species richness, Simpson’s diversity, Shannon entropy, and Pielou’s evenness, showed no significant correlations with the FAST score (Supplementary Figure S2), and PERMANOVA explained only a small fraction of community-level variation. Thus, the disease-associated microbial patterns observed here are not well described as simple increases or decreases in overall diversity. Rather, the ELA framework revealed that specific combinations of taxa formed representative community compositions associated with low, intermediate, and high FAST-score ranges. This supports the view that liver disease progression is accompanied by reorganization of community structure rather than by uniform diversity loss.

The representative community compositions further suggested a transition from low-FAST-associated states to intermediate- and high-FAST-associated states. Low-FAST compositions were characterized by taxa such as *Anaerostipes hadrus*, *Bacteroides massiliensis*, and unclassified taxa assigned to *Subdoligranulum*, *Firmicutes* and *Dorea* whereas intermediate- to high-FAST compositions were characterized by the appearance or increased importance of an unclassified *Ruminococcus gnavus* group taxon and *Streptococcus salivarius* (Fig. 3a-c). This pattern suggests that disease progression may involve not only the enrichment of taxa positively associated with FAST score, but also the loss or replacement of taxa associated with healthier community configurations. Importantly, taxa with similar environmental preferences tended to form mutually exclusive relationships, indicating that alternative community compositions can emerge even under comparable FAST-score conditions.

The community shaping index (CSI) provided an additional layer of interpretation by quantifying how strongly individual taxa were associated with each representative community structure. Overall, taxa with *g*_*FAST*_ values close to zero tended to show large variation in CSI, and no significant correlation was observed between *g*_*FAST*_ and CSI (Fig. 4). This indicates that taxa strongly associated with the FAST-score environment are not necessarily the same taxa that shape the corresponding community composition. In other words, the taxa most responsive to liver disease severity and the taxa most important for organizing microbial community structure may only partially overlap. This distinction is central to the present framework, because it separates taxon‒environment associations from taxon‒taxon relationships within a unified model.

Among the low-FAST-associated community compositions, an unclassified *Subdoligranulum* showed consistently high CSI values (Fig 4a-c) , and the exploratory FASK analysis placed this taxon upstream of both the FAST score and community composition in the inferred graphs. This result is biologically plausible, as *Subdoligranulum* species are recognized butyrate producers, and butyrate is known to support intestinal barrier function and exert anti-inflammatory effects. Consistent with these functional properties, previous studies have associated *Subdoligranulum* with metabolic health and protective gut profiles (Louis et al. 2016; Zhang et al. 2017; Cani et al. 2021), and enrichment of this genus has been reported in healthy controls compared with patients with liver cirrhosis (Liu et al. 2025). Together, these results suggest that the unclassified *Subdoligranulum* may play a potentially active role in linking healthier liver function with coherent low-FAST-associated community organization, although longitudinal or interventional validation will be required to determine whether it directly contributes to maintaining a healthy gut‒liver axis.

In contrast, the *R. gnavus* group taxon showed high CSI values in representative compositions associated with intermediate-to-high FAST scores (Fig. 4e, g, h). Interestingly, the *R. gnavus* group taxon showed a strong mutually exclusive relationship with the unclassified *Subdoligranulum* (Fig. 3g), suggesting that these taxa may occupy contrasting ecological positions in the transition from low-FAST to intermediate- or high-FAST community structures. This contrast is consistent with previous reports linking *R. gnavus* to metabolic dysregulation, inflammation, bile acid metabolism, mucin degradation, and increased intestinal permeability (Henke et al. 2019; Wu et al. 2023). However, in our causal discovery analysis, the *R. gnavus* group taxon did not consistently emerge as a strong causal driver of either liver condition or overall community structure. Further validation using larger and more multidimensional datasets will be needed to clarify its functional and causal role.*Streptococcus salivarius* represented a different pattern. This taxon showed the highest *g*_*FAST*_ value and had the second-highest CSI in C8 (Fig. 4h), the composition associated with the highest FAST scores. Moreover, the exploratory FASK analysis placed S. salivarius upstream of both the FAST score and community composition in the inferred graph. These findings are suggestive because *S. salivarius* has been repeatedly associated with more severe liver disease states. Previous studies reported increased *S. salivarius* abundance in cirrhotic patients with hepatic encephalopathy (Zhang et al. 2013), in patients with advanced fibrosis (Take et al. 2023), and in obese children with MASH (Wei et al. 2024). Because *S. salivarius* is a urease-producing bacterium (Yukawa-Muto et al. 2022), its expansion in the gut may contribute to ammonia production and pathological gut‒liver interactions. Our results extend these previous associations by suggesting that S. salivarius is linked not only to advanced liver disease, but also to high-FAST-associated dysbiotic community structures.

Several taxa with high *g*_*FAST*_ values, including oral cavity-associated anaerobes such as *Fusobacterium mortiferum*, *Dialister invisus*, and *Veillonella ratti*, should be interpreted differently. These taxa may reflect oral-to-gut translocation or opportunistic expansion under dysbiotic conditions, a phenomenon previously reported in liver disease (Qin et al., 2014, Chen et al. 2016). However, their CSI values were not consistently high in the high-FAST-associated community compositions. This suggests that some taxa may be strongly associated with the disease environment without necessarily playing a central role in shaping the characteristic community structure. Thus, integrating *g*_*FAST*_ with CSI helps distinguish environmentally responsive taxa from taxa that are more directly involved in community organization.

Our results also highlight intragenus divergence: in other words, some species within the same genus exhibited opposing associations with FAST score. For example, *B. coprocola* and *S. salivarius* were linked to high scores, whereas *B. fragilis* and *S. thermophilus* were linked to low scores. Similarly, in the genus *Fusobacterium*, *F. mortiferum* and an unidentified species showed opposite trends. These findings reinforce the need for species- or strain-level resolution in functional interpretation, rather than relying on coarse genus-level annotations. Interestingly, we also found that traditionally beneficial genera, such as butyrate-producing *Lachnospira* and *Lachnoclostridium* and lactic acid bacteria including *Lactobacillus* and *Bifidobacterium* (*B. pseudocatenuatum*), had species that were more abundant in individuals with high FAST scores. These associations were primarily driven by unclassified species, suggesting potential context-dependent roles and underscoring the need for more refined functional profiling.

Our study demonstrates the potential role of ELA to function as a bridge between microbiome data and clinical decision-making. When comparing the low-, intermediate-, and high-FAST score ranges derived from previous clinical validation with the provisional ranges inferred here from microbiota structure, we found that community compositions containing the *R. gnavus* group taxon, which characterized intermediate- to high-FAST-associated microbiota, appeared before the conventional low-FAST threshold of ε_FAST_ = 0.12 (FAST score of 0.35). This observation suggests that microbiota-based community transitions may begin to emerge before samples exceed the clinically defined low-FAST range, although further validation is needed to determine its clinical significance. These findings have potential implications for microbiome-based biomarkers. The FAST score is widely used as a noninvasive indicator of progressive MASH and liver fibrosis, but its interpretation can be influenced by host factors such as BMI, metabolic status, and other clinical covariates (Mare et al., 2024; Perazzo et al., 2021; O’Hara et al., 2024). Incorporating microbiome-derived ecological information (Ning & Hong, 2024) may improve the biological interpretation of FAST-score-associated disease states.

Another important point worth emphasizing is that our framework unifies information that is typically obtained from different methods in microbiome statistical analysis. In practice, we show that ELA can substitute for multiple conventional analyses, including supervised learning (Fig. 2), clustering (Fig. 3e), differential abundance analysis (Fig. 3f), and network inference (Fig. 3g). Notably, in interpreting the results of the latter two analyses together (Fig. 4), our framework offers a key advantage: it decomposes the effects of environmental and taxa-taxa relationships within the same model. This decomposition is further strengthened by exploratory causal analysis, which allows us to distinguish taxa that are merely correlated with environmental gradients from those that may potentially play active roles in influencing host conditions and community shifts. This integration enables a more mechanistic and interpretable understanding of microbiome dynamics than is possible by combining separate statistical approaches.

Several limitations should be noted. First, although ELA can detect multiple representative community compositions and potential state transitions, these should not automatically be interpreted as definitive evidence of true dynamical multistability. The inferred energy minima are local minima in the discrete community-membership space and should not be directly equated with stable equilibria. Such membership configurations could instead correspond to neighborhoods of stable attractors, regions near separatrices or saddle-associated bottlenecks, or coarse-grained configurations frequently visited along transient trajectories. Distinguishing among these possibilities requires time-resolved data or explicit dynamical modeling beyond the present cross-sectional ELA framework. Second, clinical metadata beyond the FAST score, such as age, BMI, weight, alcohol intake, and comorbidities, were not fully incorporated into the landscape model. Some observed heterogeneity may therefore reflect unmeasured covariates rather than intrinsic microbial state structure. Third, the causal discovery analysis should be regarded as exploratory. Although FASK can suggest possible causal directionality from cross-sectional data, longitudinal and interventional studies are required to determine whether taxa such as unclassified *Subdoligranulum* or *S. salivarius* actively drive changes in liver condition and community structure.

Fourth, he current model assumes a linear environmental effect of the FAST score on taxon occurrence, which may not capture taxa that preferentially occur at intermediate disease states or under more complex nonlinear host‒microbiome interactions. Finally, because this study was based on 16S rRNA gene sequencing, the taxonomic and functional resolution of several key unclassified taxa and species groups remains limited. Shotgun metagenomic sequencing would help resolve these taxa at species or strain level and provide functional information needed to strengthen their ecological interpretation.

Future studies should validate these findings in larger and longitudinal cohorts with richer clinical, dietary, metabolomic, and metagenomic information. Longitudinal data will be particularly important for distinguishing whether transitions in gut microbiota composition reflect within-individual liver disease progression, causal effects of microbiota structure on liver condition, or both processes occurring simultaneously. Functional profiling will also be required to clarify whether the taxa highlighted here contribute to disease progression. With these extensions, the ELA-based framework introduced here may support more personalized and mechanistically informed approaches to microbiome-based risk stratification and intervention in liver disease.

Overall, our study shows that FAST-score-associated liver disease progression is accompanied by structured changes in gut microbial community organization. By ELA based framework integrating, taxon-specific environmental responses, community shaping indices, and exploratory causal discovery, we identified candidate taxa that may link host liver condition with microbial community structure. These findings suggest that ecological modeling can provide a mechanistic and interpretable foundation for understanding microbiome transitions along clinical disease gradients.

## Author contributions

K.S. contributed to conceptualization, formal analysis, investigation, methodology, software development, validation, visualization, and writing of the original draft. Software development was further supported by S.T. Data curation was carried out by T.K. and H.F. All authors, including K.S., T.K., H.F., S.T., Y.Y.-M., N.K., S.F., H.M., and N.O., contributed to reviewing and editing the manuscript. N.O. was responsible for funding acquisition.

